# Insights into the stability of a therapeutic antibody Fab fragment by molecular dynamics and its stabilization by computational design

**DOI:** 10.1101/644369

**Authors:** Nuria Codina, Cheng Zhang, Nesrine Chakroun, Paul A. Dalby

## Abstract

Successful development of protein therapeutics depends critically on achieving stability under a range of conditions, while retaining their specific mode of action. Gaining a deeper understanding of the drivers of instability across different stress conditions, will potentially enable the engineering of protein scaffolds that are inherently manufacturable and stable. Here, we compared the structural robustness of a humanized antibody fragment (Fab) A33 using atomistic molecular dynamics simulations under two different stresses of low pH and high temperature. RMSD calculations, structural alignments and contact analysis revealed that low pH unfolding was initiated through loss of contacts at the constant domain interface (C_L_-C_H_1), prior to C_L_ domain unfolding. By contrast, thermal unfolding began with loss of contacts in both the C_L_-C_H_1 and variable domain interface (V_L_-V_H_), followed by domain unfolding of C_L_ and also of V_H_, thus revealing divergent unfolding pathways. FoldX and Rosetta both agreed that mutations at the C_L_-C_H_1 interface have the greatest potential for increasing the stability of Fab A33. Additionally, packing density calculations found these residues to be under-packed relative to other inter-domain residues. Two salt bridges were identified that possibly drive the conformational change at low pH, while at high temperature, salt bridges were lost and reformed quickly, and not always with the same partner, thus contributing to an overall destabilization. Sequence entropy analysis of existing Fab sequences revealed considerable scope for further engineering, where certain natural mutations agreed with FoldX and Rosetta predictions. Lastly, the unfolding events at the two stress conditions exposed different predicted aggregation-prone regions (APR), which would potentially lead to different aggregation mechanisms. Overall, our results identified the early stages of unfolding and stability-limiting regions of Fab A33, which provide interesting targets for future protein engineering work aimed at stabilizing to both thermal and pH-stresses simultaneously.

**Author Summary:** Currently, antibody-based products are the most rapidly growing class of pharmaceuticals due to their high specificity towards their targets, such as biomarkers on the surface of cancer cells. However, they tend to aggregate at all stages of product development, which leads to decreased efficiency and could elicit an immunological response. Improvements in the stability of therapeutic antibodies are generally made during the development phase, by trial and error of the composition of the formulated product, which is both costly and time consuming. There is great demand and potential for identifying the drivers of instability across different stress conditions, early in the discovery phase, which will enable the rational engineering of protein scaffolds. This work elucidated the stability-limiting regions of the antibody fragment Fab A33 using several computational tools: atomistic molecular dynamics simulations, *in-silico* mutational analysis by FoldX and Rosetta, packing density calculators, analysis of existing Fab sequences and predictors of aggregation-prone regions. Results identified particular regions in which mutagenesis has the potential to stabilize Fab against both thermal and pH-stresses simultaneously. Overall, the methodology used here could improve the developability screening of candidate antibody products for many diseases, such as cancer, chronic inflammatory diseases and infectious diseases.

## Introduction

In the last 30 years, monoclonal antibody products have become the main drug class for new approvals in the pharmaceutical industry [1]. To date, over 60 antibody-based drugs are on the market, representing half of the total sales, with over 550 further antibodies in clinical development [2]. They are used as therapeutic drugs to treat human diseases, mainly in oncology, auto-immune diseases and cardiovascular diseases. The use of antibody fragments, such as the antigen-binding antibody fragment (Fab) studied here, brings additional advantages, including deeper tissue penetration due to their smaller size, which has proven beneficial to treat tumors [3]. In addition, Fab fragments lack the Fc domain, and thus are not glycosylated which allows simpler and less costly manufacture due to their expression in prokaryotic systems [4]. However, the lack of the Fc domain leads to their more rapid clearance in humans than for full antibodies.

The stabilization of therapeutic proteins against aggregation remains one of the biggest challenges facing approvals as biopharmaceutical products [5–7]. Thus, not only their mode of action, but protein stability is a crucial factor to their becoming successful products. Novel antibody products such as Fabs, single-chain variable fragments (ScFvs) and bi-specifics are currently being developed and their properties remain largely unknown. Knowledge about the stability of these pharmaceutical products, especially in the early development stages, would aid in their engineering and the design of antibody fragments that are more aggregation resistant.

Native protein conformations are only marginally stable, and are highly dynamic, hence they are more realistically described as a native ensemble. There is increasing evidence to suggest that under native conditions, aggregation takes place primarily from partially unfolded native-like states [8–12]. However, little is known about the structures of native conformers that initiate aggregation, or how these are affected by different stress conditions. Local unfolding of proteins can expose aggregation prone-regions (APR), that have the potential to trigger aggregation [13,14]. APRs are the regions in the protein most likely to form and stabilize the cross-β structures that are characteristic of many aggregates, notably hydrophobic sequences with low net charge and a strong β-sheet forming propensity. Generally, APRs are located in the protein core, protected from the solvent, and thus blocked from forming cross-β structure. Under certain stresses, such as an increase in temperature, a change in pH, addition of denat
urants, or elevated shear force, structural regions in the protein may destabilize and partially unfold to expose any underlying APRs [15]. Each structural region of the protein can respond differently to stress, and so the overall pattern of responses is likely to vary with each type of stress. Thus, determining the conformational changes that a protein experiences under different stress conditions is important for identifying common routes towards stabilization across all stress conditions via either mutagenesis or formulation.

Molecular dynamics (MD) simulations have been extensively used to study protein stability [16–21]. MD simulations offer atomic resolution insights into the early conformational events that can take place under different conditions. To date, not many all-atom MD studies on antibody fragments have been reported. MD simulations were used previously to study the aggregation potential of an antibody Fab fragment, from a human IgG1k antibody, via multiple elevated temperature MD simulations at 300 K, 450 K and 500 K [22]. This revealed that domain interfaces deformed prior to the unfolding of individual domains, and that two V_H_ domain sites were potentially labile to aggregation. Their structural deformation increased the solvent-accessible surface area of the APRs in those regions. The unfolding process of an antibody Fab fragment was also studied using an elastic network model, to reveal that the constant regions were more flexible, and unfolded earlier, than the variable regions [23]. MD simulations at 450 K and 500 K have also revealed the stability-limiting regions of an antibody single-chain variable fragment (scFv) [24]. Disruption of the V_L_-V_H_ interface was found to precede the unfolding of the domain structures. In contrast to the study on the Fab above, the V_H_ domain of the scFv was found to be more thermally resistant than the V_L_ domain.

Each Fab is composed of one light and one heavy chain (Fig 1 and Fig S1), each comprising a variable (V_L_ or V_H_) and a constant (C_L_ or C_H_1) domain. Each domain forms an immunoglobulin fold, having two layers of β-sheets, an inner β-sheet and an outer β-sheet. Constant domains are formed of seven β-strands in a 4+3 arrangement, while variable domains have two additional β-strands in a 5+4 arrangement. The variable domains contain the antigen-binding site at their complementary determining regions (CDRs), formed by three loops in V_L_ and three loops in C_L_. There are five disulfide bonds in Fab, four of them intra-domain and the last one between the light and heavy chains at the hinge region. Individual domains interact to form the variable region interface (V_L_-V_H_), and the constant region interface (C_L_-C_H_1). Interface contacts are shown in Fig 1 and the residues involved in the contacts are listed in Table S1. The variable region interface is mainly formed by aromatic side chains that are tightly packed and located at the center of the interface (six Tyr, two Trp, and two Phe), forming hydrophobic interactions. However, fewer aromatic side-chains are involved in the constant region domain interface (four Phe). Furthermore, no contacts at <3.5 Å were found between part of one of the β-strands (at residues 177-180) in the C_L_ domain, and the C_H_ domain in our Fab A33 homology model (Fig 1).

**Fig 1.**
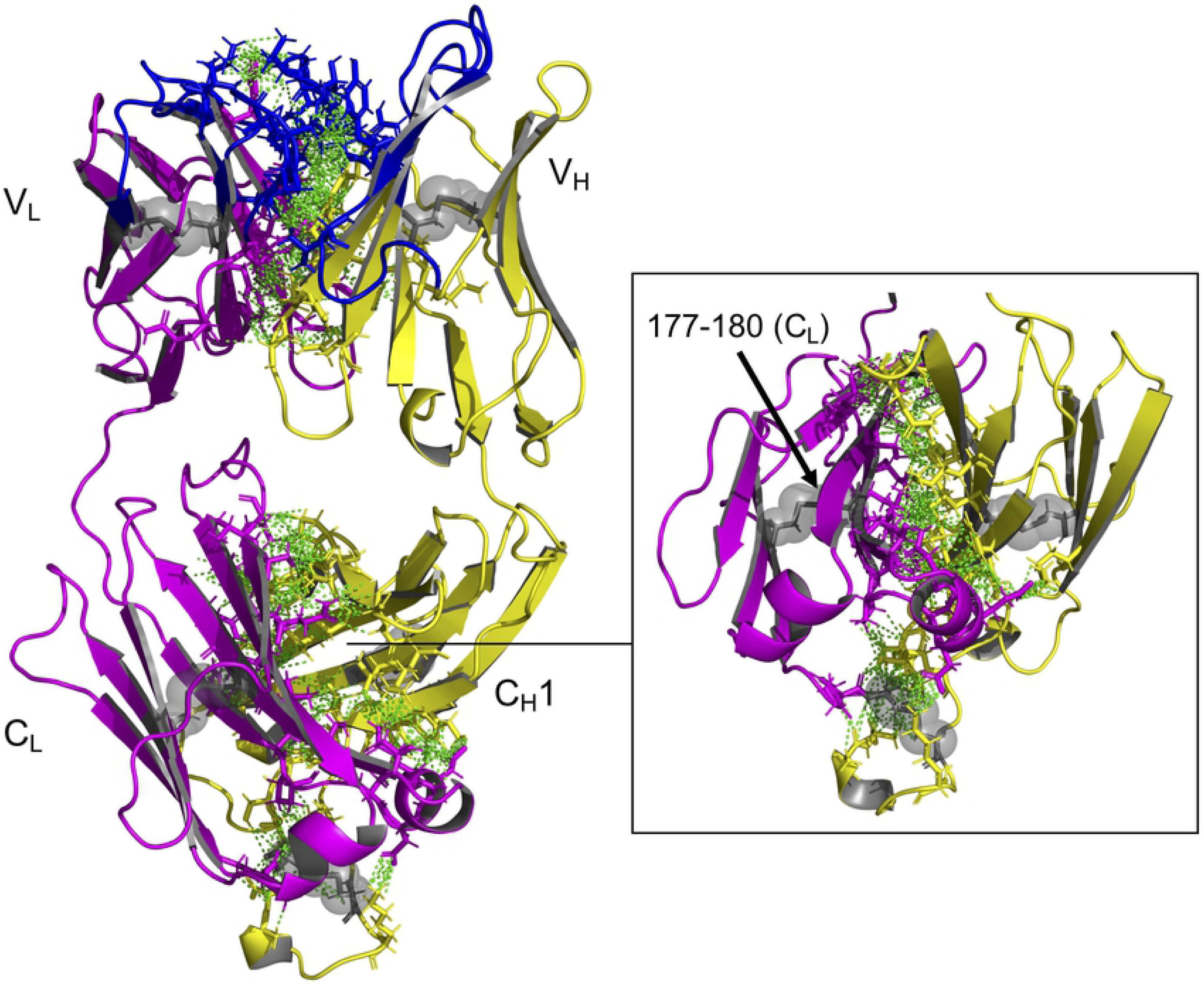
Fab A33 structure with interface contacts highlighted. Fab is composed of light (magenta) and heavy (yellow) chains. Each chain contains variable (V_L_ and V_H_) and constant (C_L_ and C_H_1) domains. The antigen-binding region at the complementary determining regions (CDRs; blue), are located in the variable domains. There are five disulfide bonds (gray highlights). Contacts between heavy and light chains within 3.5 Å are indicated with green dashed arrows. Β -strand 177-180 in C_L_ domain does not have contacts with C_H_ domain, zoom in right-inset.

Here, we compare the early unfolding events of Fab A33 at both high temperature and low pH, using all-atom MD simulations. A common feature to both stress conditions was that unfolding was initiated by the loss of interfacial contacts between neighboring domains, and that domain unfolding occurred later. However, our results revealed different unfolding pathways for the two stress conditions, leading to partial unfolding of only the C_L_ domain at low pH, compared to destabilization of both C_L_ and V_H_ domains in the high temperature condition. These conformational changes exposed different predicted aggregation-prone regions (APR), which would additionally support divergent aggregation mechanisms. Salt-bridge analysis provided insights into the location of those that were broken most rapidly due to protonation in the low pH simulation, and also showed that high temperature led to an increased fluctuation of salt bridge formation and breaking, more generally throughout the structure. An *in-silico* mutational analysis by both FoldX and Rosetta, predicted that the constant domain interface had the greatest potential for further stabilization, a finding that was also supported by lower packing-density calculations. Taken together, our results determined the stability-limiting regions at low pH and high temperature for Fab A33, and also identified those with the greatest potential for mutations that simultaneously improve stability under both low pH and high temperature conditions.

## Results

### Interface contacts, RMSD of individual domains and structural alignments revealed different unfolding pathways at low pH and high temperature

To determine which domains of Fab A33 are more susceptible to unfolding under low pH and high temperature, we first followed the RMSD of each individual domain (V_L_, V_H_, C_L_ and C_H_1) along the simulations, as changes in RMSD are indicative of a conformational change. Simulations in the unfolding trajectories (pH 3.5 and pH 4.5 at 300 K, for low pH; pH 7.0 at 340 K and 380 K, for high temperature) were compared to the simulations in the native trajectory (pH 7.0 at 300 K). For every condition of pH and temperature, three independent simulations were performed, and their average RMSD are shown in Figures 2 and 3. Additionally, structures from the unfolding trajectories (pH 3.5, for low pH; 380 K, for high temperature) were aligned to structures from the native trajectory (pH 7.0 at 300 K), to visualize the structural changes that individual domains experienced. For each domain alignment, ten structures were taken every 3 ns from each simulation repeat, from the 20-50 ns range at which the RMSD had stabilized. Thus, a total of thirty structures from each stress condition were compared to thirty from the native trajectory. We also monitored the number of interface contacts between domains (V_L_-V_H_ and C_L_-C_H_1) during the simulations using a cutoff of 4 Å, to understand the temporal relationship between breakage of contacts in each interface, and the unfolding of each domain. The RMSD and radius of gyration (R_g_) of the whole protein are also shown at every condition in Fig S2; where increased RMSD and R_g_ were observed at the conditions of low pH (3.5) and high temperature (380 K).

**Fig 2.**
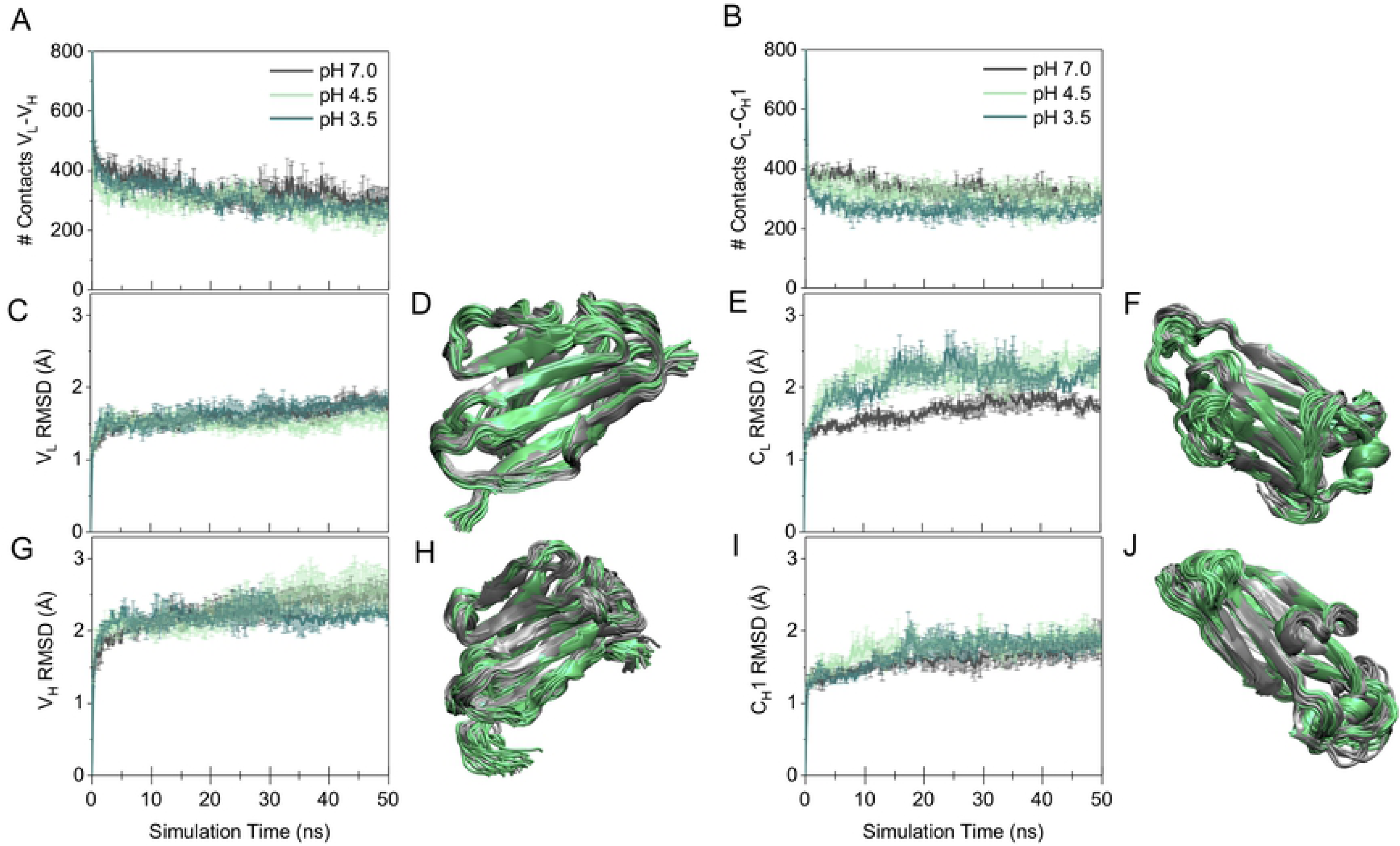
Interface contacts, RMSD of individual domains and structural alignments for simulations at pH 7.0, 4.5 and 3.5 (all 300 K). A, B) Contacts between light and heavy chains within 4.0 Å with simulation time, for variable (V_L_-V_H_) and constant (C_L_-C_H_) regions, respectively, pH values as labelled. C, E, G, I) RMSD of individual domains with simulation time for V_L_, V_H_, C_L_ and C_H_1, respectively, pH values as labelled. In all cases, the average of three independent simulations is shown with the SEM as error. D, F, H, J) Alignments of structures from simulations at pH 7.0 and 3.5 for V_L_, V_H_, C_L_ and C_H_1, respectively. Ten structures from the last 30 ns of each simulation were used, totaling thirty structures from pH 7.0 and thirty from pH 3.5.

**Fig 3.**
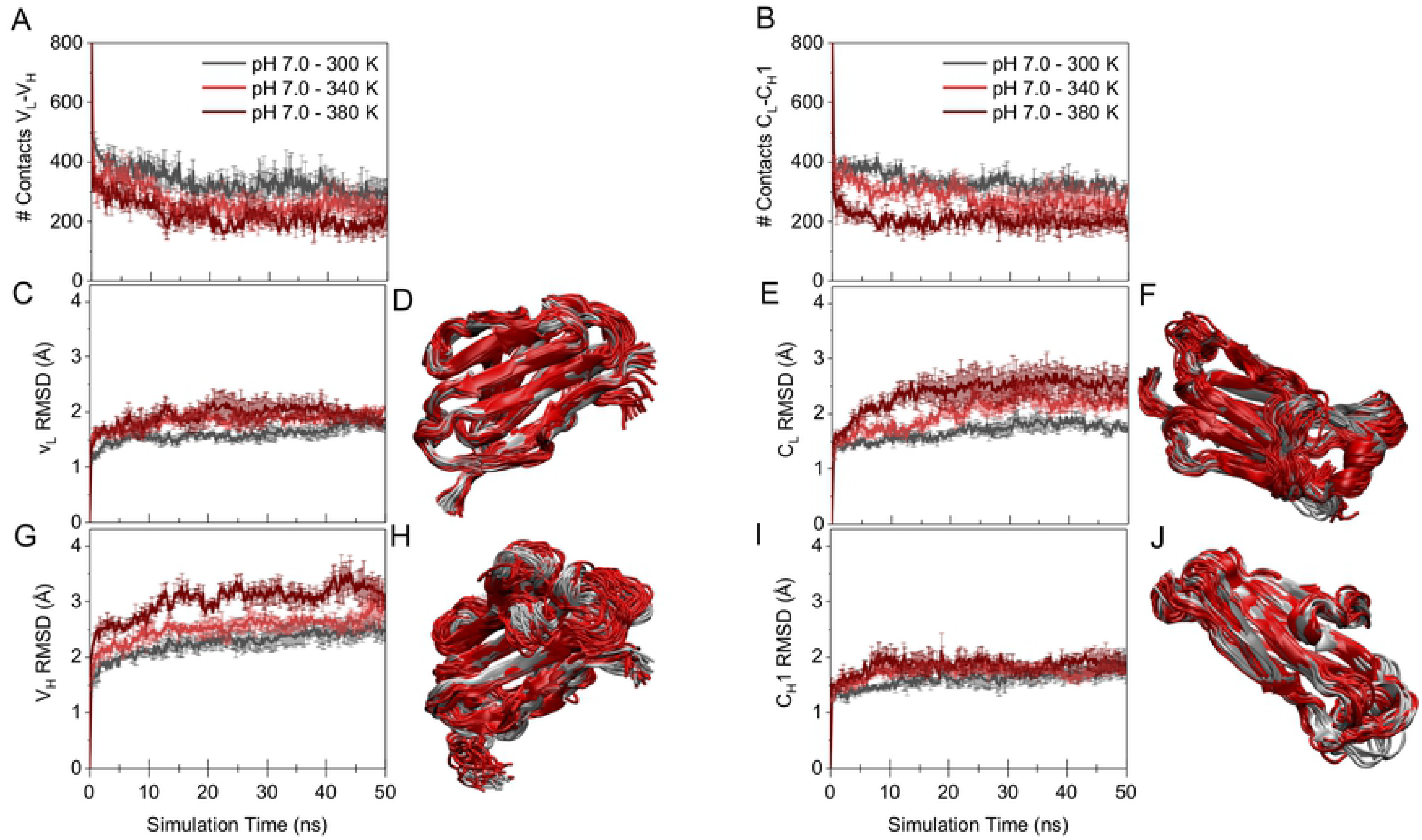
Interface contacts, RMSD of individual domains and structural alignments for simulations at pH 7.0 and temperatures 300 K, 340 K and 380 K. A, B) Contacts between light and heavy chains within 4.0 Å with simulation time, for variable (V_L_-V_H_) and constant (C_L_-C_H_) regions, respectively, temperature values as labelled. C, E, G, I) RMSD of individual domains with simulation time for V_L_, V_H_, C_L_ and C_H_1, respectively, temperature values as labelled. In all cases, the average of three independent simulations is shown with the SEM as error. D, F, H, J) Alignments of structures from simulations at temperatures of 300 K and 380 K for V_L_, V_H_, C_L_ and C_H_1, respectively. Ten structures from the last 30 ns of each simulation were used, totaling thirty structures from 300 K and thirty from 380 K.

At low pH, almost no change was observed in the number of interfacial contacts within the variable region (V_L_-V_H_) of Fab A33, when compared to at pH 7.0. For pH 7.0, 333 ± 24 contacts were maintained (discarding the first frame), and 309 ± 24 contacts were maintained at pH 3.5 (Fig 2A). By contrast, a loss of interfacial contacts in the constant region (C_L_-C_H_1) was observed between pH 7.0 with 335 ± 17 contacts, and pH 3.5 with 265 ± 12 contacts (Fig 2B). Interestingly, this loss of constant region interfacial contacts at low pH took place very quickly, with pH 7.0 retaining 384 ± 14 contacts after 5 ns of the simulation, while simulations at pH 3.5 retained only 270 ± 11 contacts. This could be attributed to the lack of a well-defined hydrophobic core in the C_L_-C_H_1 interface, resulting in numerous early-disrupted contacts. Notably, C_L_ was the only domain to show a noticeable conformational change at low pH, revealed as an increase in RMSD from 1.8 ± 0.1 Å at pH 7.0 (calculated between 20-50 ns of the simulation), to 2.2 ± 0.1 Å at pH 3.5 (Fig 2E). This domain displacement occurred in the first 20 ns, after many interface contacts had already been lost with respect to pH 7.0, which suggests that destabilization of the C_L_-C_H_1 interface preceded and potentially accelerated the unfolding of the C_L_ domain. The other domains (V_L_, V_H_ and C_H_1) did not unfold significantly during the low pH simulations (Fig 2C, 2G and 2I). Structural alignments confirmed this result, showing remarkably good alignment between the pH 7.0 and pH 3.5 structures in each case for V_L_, V_H_ and C_H_1 (Fig 2D, 2H and 2J). Alignments of the C_L_ domain at pH 7.0 and pH 3.5 revealed a slight displacement at low pH, especially visible in the loop regions (Fig 2F). These findings agreed with previous experimental work, which combined SAXS, atomistic modelling and smFRET to reveal the displacement of the C_L_ domain in Fab A33 at low pH [14].

For thermal denaturation, MD simulations are commonly run at temperatures as high as 500 K to attempt to fully denature the protein. Here, we aimed to capture the early thermal unfolding events of Fab A33, which involve only partial unfolding of the protein. For this reason, and to reflect experimental conditions more closely, lower temperatures of 340 K and 380 K were used in our simulations. Interfacial contacts were found to break across both the variable and the constant regions (Fig 3A and 3B). At 380 K, there was an average of only 220 ± 24 contacts in the variable interface and 204 ± 13 contacts in the constant interface. Those between the constant domains broke earlier than those of the variable domains, with only 218 ± 14 present after just 5 ns of the simulations at 380 K. This is consistent with previous reports, which also found that the constant region interface lost a larger fraction of its total interface contacts consistent
ly faster than the variable region interface at high temperature, and also that domain unfolding occurred later than the loss of interfacial contacts [22]. Overall, more contacts were lost at both interfaces with high temperature than at low pH.

While V_L_ and C_H_1 experienced only small domain displacements (Fig 3C and 3I), clear domain unfolding was observed for both C_L_ and V_H_ (Fig 3E and 3G). At 380 K from 20 ns to 50 ns, the V_H_ domain displayed an increase in RMSD from 2.4 Å at pH 7.0, to 3.2 Å at pH 3.5. That for the C_L_ domain increased from 1.8 Å at pH 7.0 to 2.4 Å at pH 3.5 (all averages were ± 0.1 Å). In these cases, many interface contacts were also lost with respect to pH 7.0 and 300 K, before the unfolding of individual structural domains, again consistent with destabilization of the interface contributing to the loss of stability for the individual domains. For both V_L_ and C_H_1, structures from the simulations at 300 K and 380 K aligned well (Fig 3D and 3J), whereas for V_H_ and C_L_ (Fig 3H and 3F) the domains were structurally perturbed at the higher temperature. The V_H_ domain experienced a displacement of the loops on the N-terminal region, including the three CDR loops (Fig 3H). Differences in the C_L_ domain at high temperature were found in the loops and within an internal β-strand (Fig 3F). This was consistent with previous work which identified instability and structural changes in the V_H_ domain of another Fab at high temperature [22]. Taken together, these findings suggest a different unfolding pathway for Fab A33 at low pH and at high temperature.

### Loss in β-strand secondary structure confirms regions of unfolding

The unfolding of individual domains was additionally followed by their loss in secondary structure (SS); specifically, we monitored the change in β-strand structure. Constant domains are composed of seven β-strands named A to G, while variable domains contain two more strands, a total of nine, with the two additional strands termed C’ and C’’ (Fig 4A and 4B). To calculate the loss in β-strand structure for each of the strands, we first tracked the secondary structure designation for each residue in Fab A33 throughout the simulations (Fig S3). The percentage of time occupied within β-strand was calculated for each residue, and then summed for each of the 32 β-strands in Fab A33. This value was averaged for each of the three repeats at each condition. The percentage change in β-strand occupancy was then calculated, to determine the loss relative to the reference simulations at pH 7, 300 K (Fig 4 and Fig S4).

**Fig 4.**
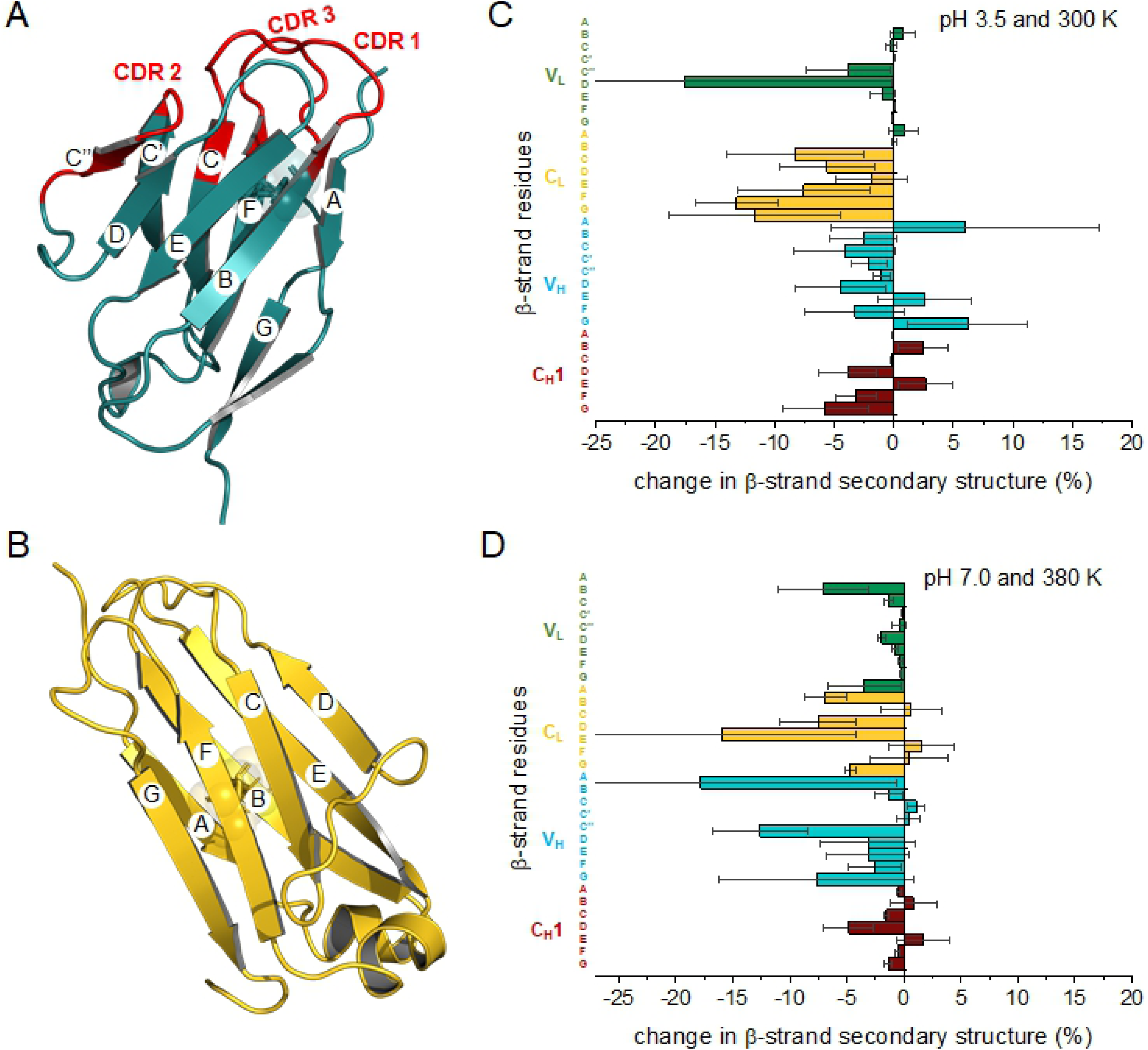
Loss of secondary structure for each of the 32 β-strands of Fab A33. A, B) Strand order shown by lettering (A-G) for variable and constant domains, respectively. C, D) Percentage increase/decrease in β-strand secondary structure for each strand in Fab during the simulations, respect to pH 7.0 and 300 K, for: C) pH 3.5 and 300 K, D) pH 7.0 and 380 K. Error bars are the same and equal for positive and negative values.

At pH 3.5, the C_L_ domain had an overall loss in secondary structure content, confirming the results found in the previous section (Fig 4C). Strands F (−13 ± 3 %) and G (−12 ± 7 %) of the C_L_ domain showed the highest β-sheet structure loss, both located in the outer β-sheet. Strands B (−8 ± 6 %), C (−6 ± 4 %) and E (−8 ± 6 %) also experienced significant losses. Additionally, the β-strand C’’ of the V_L_ domain also showed a large variability between repeat trajectories. C’’ is the shortest strand, and is located at the extreme of the outer β-sheet connecting the CDR-2 loop, and so the large variability suggests that this is a flexible region prone to the loss of secondary structure in some but not all simulations.

At 380 K, the losses in β-strand content were located in both the C_L_ and V_H_ domains, consistent with the unfolding described in the previous section (Fig 4D). Many strands in the C_L_ domain showed significant losses, A (−7 ± 2 %), C (−8 ± 3 %), D (−16 ± 12 %) and G (−5 ± 1 %), located at the extremes of the inner and outer β-sheets. The V_H_ domain also showed high losses of β-strand. However, of these strands A (−18 ± 17 %) and G (−8 ± 9 %) also showed high variability between repeats. Interestingly, these same two strands in V_H_ were previously found to deform at high temperature in a different Fab [22]. Strand C’’ (−13 ± 4 %) of the V_H_ domain also showed a significant loss of β-strand content.

### Salt bridge analysis identifies key stabilizing salt bridges

To identify the ionizable residues that potentially drive the conformational changes at low pH, a salt bridge analysis was performed. Salt bridges were identified over the simulation time for all the MD simulations carried out, using an O-N bond distance cutoff of 3.2 Å. From these, the occurrence (%) of each salt bridge during the simulation was calculated, and averaged for the three independent repeats at each condition. Lastly, the most persistent salt bridges at each condition were highlighted in the Fab A33 structure (Fig 5 and Fig S5).

**Fig 5.**
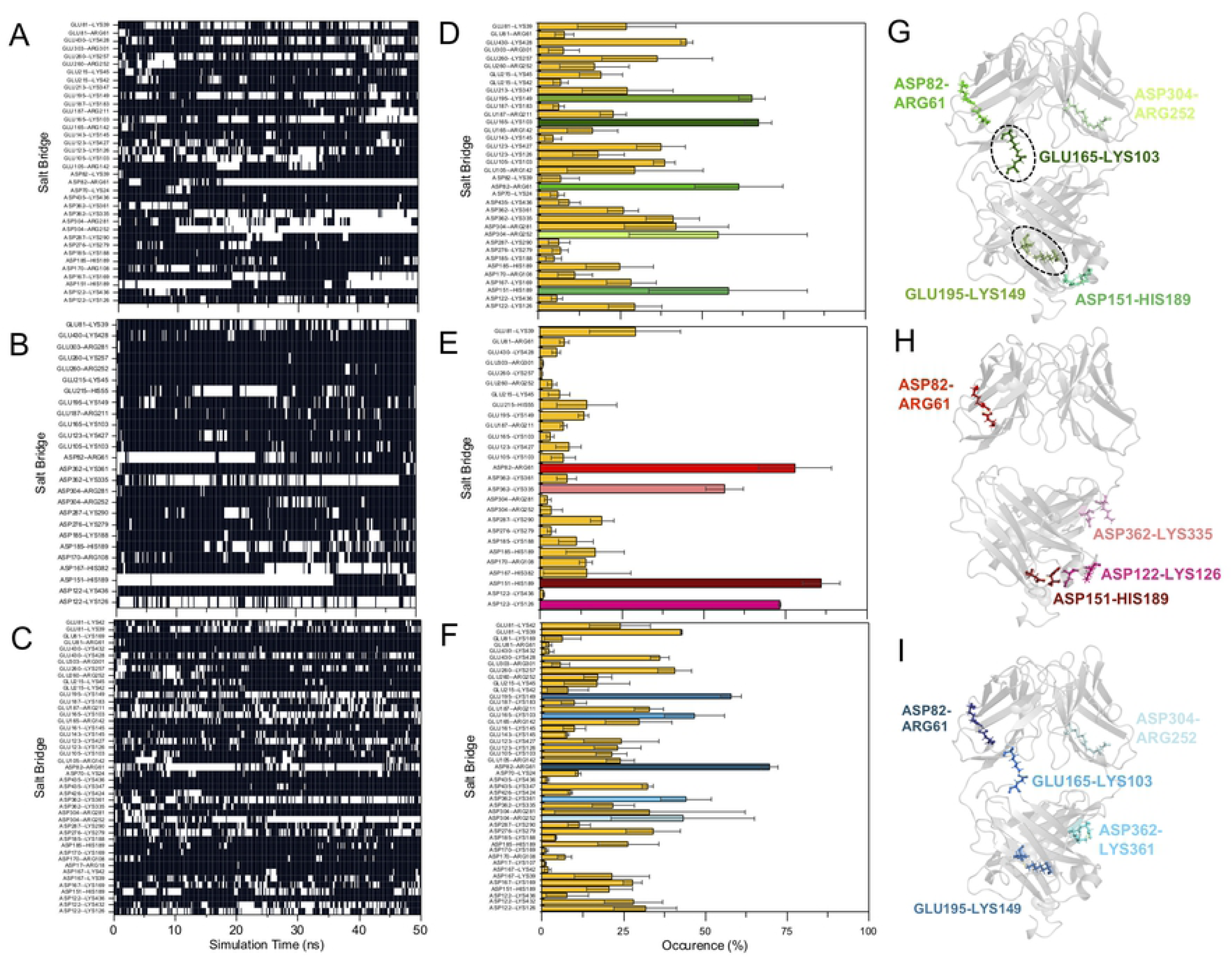
Salt bridge analysis. A, B, C) Salt bridges formed during the simulation time for representative MD simulations at A) pH 7.0 and 300 K, B) pH 3.5 and 300 K and C) pH 7.0 and 380 K. Presence of a salt bridge is indicated in white and absence in black. D, E, F) List of salt bridges and its occurrence for simulations at D) pH 7.0 and 300 K, E) pH 3.5 and 300 K and F) pH 7.0 and 380 K. Values shown are the average of three independent simulations with their SEM as error. The more persistent salt bridges are highlighted for pH 7.0 (green), pH 3.5 (red) and pH 7.0 and 380 K (blue). G, H, I) The more persistent salt bridges are mapped into the Fab A33 structure. Two key stabilising salt bridges (Glu165-Lys103 and Glu195-Lys149) are highlighted in a dashed circle.

At pH 7.0 and 300 K, a total of 36 salt bridges were present. Interestingly, many of these salt bridges were flexible and able to form with different partners during a single trajectory, such as Asp122 which partnered with both Lys126 and Lys436, or Asp304 which paired with both Arg252 and Arg281. This is consistent with previous work, which found that salt bridges break and reform, and not always with the same partner [25]. The most persistent (as % of time present) salt bridges at pH 7.0 were Glu165-Lys103 (67 ± 4 %), Glu195-Lys149 (65 ± 4 %), Asp82-Arg61 (61 ± 14 %), Asp151-His189 (58 ± 24 %), and Asp304-Arg252 (55 ± 27 %), as shown in Fig 5A, 5D and 5G.

At low pH, pH 3.5, 300 K, a total of 27 salt bridges were observed, but most of them were very short lived. The more persistent salt bridges at pH 3.5 were Asp151-His189 (86 ± 6 %), Asp82-Arg61 (78 ± 11 %), Asp122-Lys126 (73 ± 1 %), Asp362-Lys335 (56 ± 6 %) (Fig 5B, 5E and 5H). The protonation state at the end of the pH 3.5 simulations was calculated again using these Fab conformations, which revealed these salt bridges to be still present due to predicted pK_a_ values of below 3.5 for these aspartates. Comparison of the salt bridges at pH 7.0 and 3.5, indicated the presence of two salt bridges that potentially trigger the conformational change observed at low pH, and thus the loss of Fab A33 stability. Glu165-Lys103 and Glu195-Lys149, were the most persistent contacts at pH 7.0, but were not present at pH 3.5. Glu165-Lys103 bridges the C_L_ domain to the V_L_ domain, and Glu195-Lys149 bridges the outer β-strands C and F of the C_L_ domain. Loss of these salt bridges at low pH, would therefore destabilize the C_L_ domain, and promote the observed C_L_ domain displacement.

At high temperature, pH 7.0 and 380 K, a total of 45 salt bridges were observed. The greater number than at 300 K, reflects an increased conformational flexibility of many salt bridges at the higher temperature, in which they often broke, but then reformed with a different partner. Indeed, at the high temperature, salt bridges broke and reformed much faster (Fig 5C). At 380 K, the total time present for the most persistent salt bridges observed at 300 K, had decreased to 47 ± 9 % for Glu165–Lys103, 58 ± 3 % for Glu195-Lys149, 21 ± 7 % for Asp151-His189, and 43 ± 22 % for Asp304-Arg252. However, Asp82-Arg61 increased in occurrence to 70 ± 3 % (Fig 5C, 5F and 5I). These findings indicate that the increased dynamics at high temperature, results in constant rupture and formation of salt bridges, and this transient disruption leaves Fab A33 more susceptible to unfolding.

### FoldX, Rosetta and packing density calculations predict sub-optimal stability of C_L_ and the C_L_-C_H_1 interface

Computational tools such as FoldX and Rosetta-ddG [26,27] predict the relative changes in folding free energy (ΔΔ*G*) between the Gibbs free energies (Δ*G*) of the protein carrying a simulated point mutation and the wild-type protein, to find those mutations that will most significantly reduce the free energy of the protein. These approaches are often also combined to find consensus predictions [28,29]. To predict stabilizing mutations in Fab A33, we calculated the ΔΔ*G* with both FoldX and Rosetta-ddG, of all possible single-mutant variants when accessing all 19 other substitutions across the 442 residue positions in Fab A33, totaling 8398 mutations. FoldX identified 1879 of these mutations as stabilizing (22.4 %), while Rosetta-ddG identified 2386 (28.4 %). Of these, 956 (11.4 %) were predicted by both algorithms. Fig S6 shows the correlation between the mutations predicted by FoldX and Rosetta, and Table S2 lists the 25 most stabilizing mutations predicted by both algorithms, with their respective ΔΔ*G* values.

Fig 6A compares the greatest stabilization predicted by FoldX and Rosetta, at each of the 442 residues in Fab A33, regardless of the specific mutation selected by each algorithm. Six residues were highlighted in magenta in Fig 6A and 6B, as those predicted by both algorithms to have the greatest potential for stabilization. These residues corresponded to S176, N137, S397, S159, S12 and T180, and all of their mutations predicted to be most stabilizing, were to more hydrophobic amino acids, such as Trp, Leu, Ile, Phe and Tyr (Fig S6 and Table S2). Four of the six residues (S176, N137, S397 and T180), are located in the constant domain interface, between C_L_ and C_H_1 domains (Fig 6B), suggesting significant potential for further stabilization within the C_L_-C_H_1 interface. Furthermore, S159 is in the C_L_ domain, but interacts with an outer β-strands of C_L_, and S12 is in the V_L_ domain, but interacts with the C_L_ domain (Fig 6B). Thus overall, the C_L_ domain has a relatively high potential for stabilization, through repacking of the C_L_-C_H_1 interface, within the C_L_ domain itself, and also through improved interaction between C_L_ and V_L_. This is consistent with the MD simulations which found the displacement of C_L_ away from the interface with C_H_, and subsequent unfolding of the C_L_ domain, to be critical steps in early or partial unfolding.

**Fig 6.**
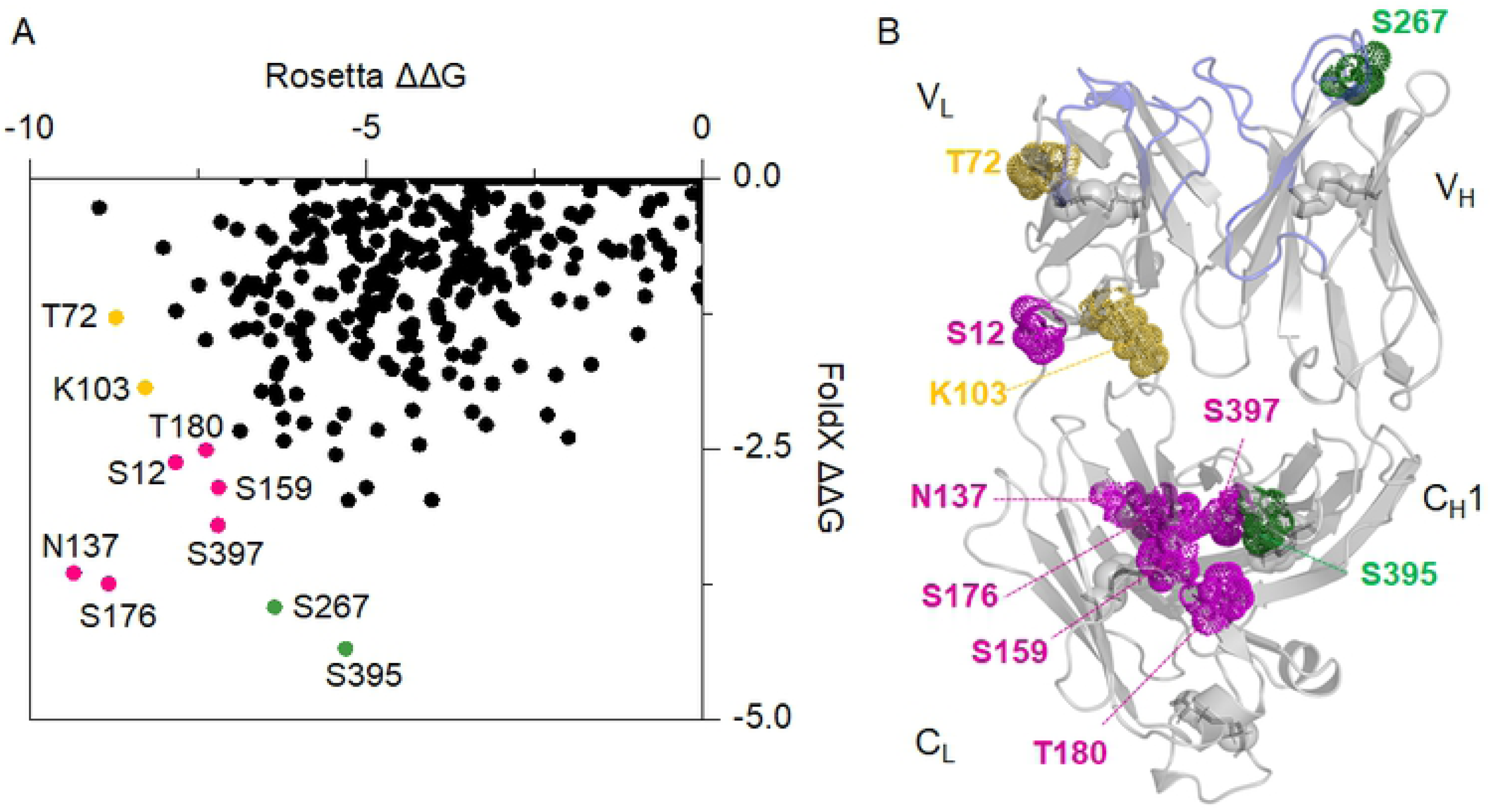
Predicted residues that can be stabilized further by FoldX and Rosetta-ddG. A) Correlation between FoldX and Rosetta predictions. Residues predicted by both software to be most stabilizing are shown in magenta on the bottom left. Residues predicted only by FoldX to be stabilizing are shown in green and mutations predicted only by Rosetta in yellow. B) Residues predicted to be stabilized further the most are mapped in Fab A33 structure, following the same colour scheme as in A).

Further highly stabilizing mutations at residues S395 and S267, as predicted only by FoldX, are highlighted in green in Fig 6. S395 is also located in the C_L_-C_H_1 interface. However, S267 is in CDR2 of the V_H_ domain, and so not a good candidate for general framework stabilization due to its role in antigen binding. Highly stabilizing mutations at residues K103 and T72, as predicted only by Rosetta-ddG are highlighted in yellow. These were both located in the V_L_ domain, but K103 also interacts with the C_L_ domain, further suggesting that the interactions within and around the C_L_ domain are the least optimized for stability.

The packing density of each Fab A33 residue was calculated using the package occluded surface (OS) software, which calculates occluded surface and atomic packing [30,31]. The occluded surface packing (OSP) value of each atom is calculated from normal vectors that extend outward from the atom surface until they intersect a neighboring van der Waals surface (Fig S7). This value is 0.0 for completely exposed residues and 1.0 where 100% of molecular surface is in contact with other van der Waals surface. The average OSP for all 28 β-strand residues within domain interfaces (V_L_-V_H_ and C_L_-C_H_1) was 0.49 ± 0.01 (OSP values shown in Table S3). By contrast, the average OSP of the five constant-domain interface residues (S176, N137, S397, T180, and S395), identified by FoldX and Rosetta as having high stabilization potential, was 0.41 ± 0.05 (OSP values shown in Table S3). This can be visualized in Fig 7, where OSP values were mapped in the structure of Fab A33, with red to indicate high packing density, and blue to indicate low packing density. β-strand residues within domain interfaces were highlighted as sticks (Fig 7A). Residues in the constant interface (C_L_-C_H_1) were lighter colored than residues in the variable interface (V_L_-V_H_), indicating less tight packing of the constant interface. An insight of the residues identified by FoldX and Rosetta is provided in Fig 7B, where a lighter color than the residues in the variable interface was also observed. This result shows that the predicted residues are under-packed, and therefore have the potential to be mutated to pack the C_L_-C_H_1 interface more tightly.

**Fig 7.**
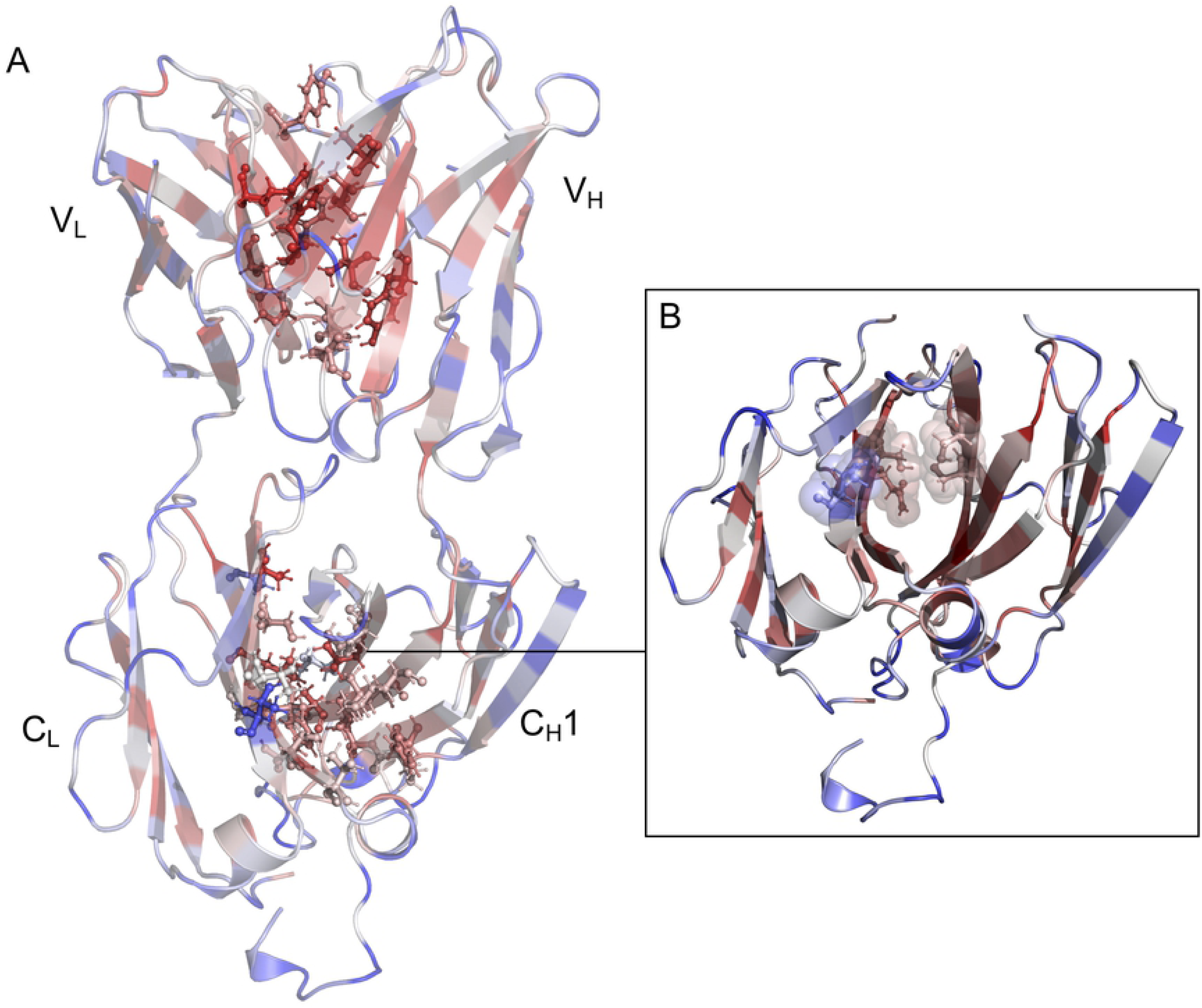
Packing density of every residue in Fab A33, computed using Occluded Surface. A) The occluded surface packing (OSP) values were added as B-factors to the PDB file for the Fab A33 homology model. High packing values are shown in red and low values in blue. Residues in β-strands within domain interfaces (V_L_-V_H_ and C_L_-C_H_1) are highlighted in sticks and ball representation. B) Residues identified by FoldX and Rosetta that could be stabilised further (S176, N137, S397, T180, and S395) are highlighted in sticks and ball, and sphere representation.

### Comparison to natural sequence variations in Fabs

The natural variability of Fab sequences was identified from within one hundred light chains and one hundred heavy chains curated from those available in the Protein Data Bank [32]. Sequence alignment and sequence entropy calculations for each residue were obtained using Bioedit [33]. An entropy of zero indicates a fully conserved residue, whereas 3.04 is the maximum entropy, originating from 21 possibilities (all amino acids plus the stop codon). There is significant positional bias in the sequence variability of Fabs due to the hypervariability of the CDRs [34], the presence of kappa (κ) and lambda (λ) light chain isotypes, allotypic diversity across individuals, and idiotypic variability within the variable domains of individuals. The sequence entropy analysis (Fig S8) clearly shows this, with the highest sequence entropy (>2), for the six CDRs, and a slightly lower variability on average within the C_H_ domain. Similarly, the higher sequence entropy on average for the C_L_ domain, compared to the C_H_ domain results from the grouping of kappa (κ) and lambda (λ) light chain isotypes.

Even though residues in the constant regions have more restricted variability, many natural variations are still observed that may affect stability. Except for the fully conserved S176, the sites predicted as having the most potential for stabilizing mutations by both FoldX and Rosetta, had natural variations, with sequence entropy values of N137: 0.74, S397: 0.15, S159: 0.96, S12: 0.96, T180: 0.43. This was also true for mutations identified only by FoldX (S267: 1.69 and S395: 0.69), and only by Rosetta-ddG (K103: 0.16, T72: 0.91). Comparisons between the existing mutations and the stabilizing mutations suggested by FoldX and Rosetta are shown in Table 1. In general, the mutations found in existing Fabs were conservative changes to residues with properties similar to the original residue. For instance, S397 and S395 are only naturally mutated to Thr, whereas T180 can also be Ser, and K103 can be Arg. By contrast, FoldX and Rosetta predictions were typically from polar to more hydrophobic residues, typically to Ile, Leu, Trp, Val, and Tyr. A few suggested mutations were also found naturally, such as N137I, S12Y, and S267P. S159 also shows the potential to be mutated to more hydrophobic residues, Val and Met. Overall, this analysis shows that despite their low natural variability, many residues in the constant domains had significant scope for stabilization through non-natural mutations.

**Table 1.**
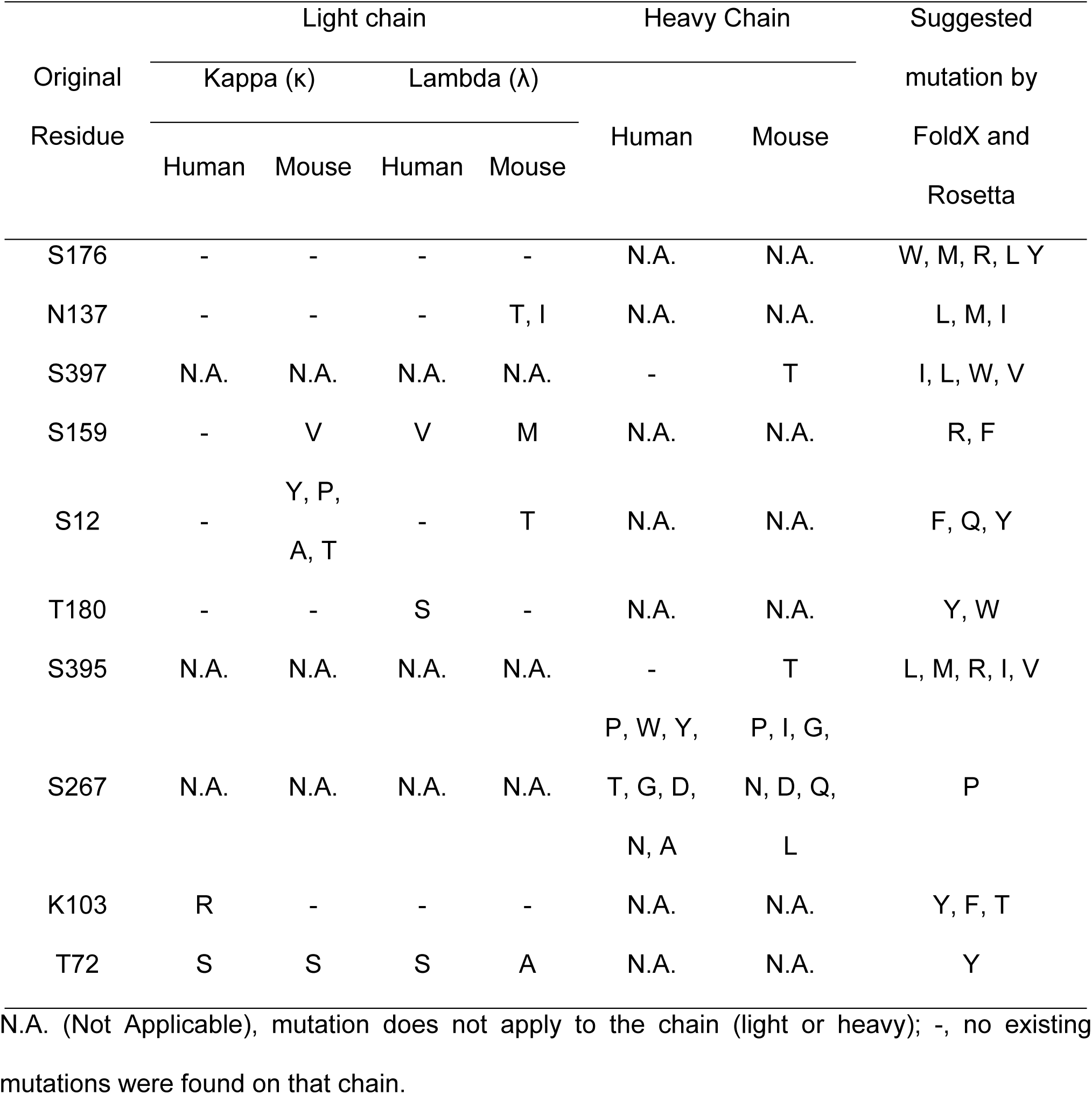
Comparison between the mutations in existing human and mouse Fabs and the stabilizing mutations suggested by FoldX and Rosetta.

### Solvent exposure of different aggregation-prone regions promotes different aggregation pathways for low pH and high temperature

The aggregation pathways of Fab A33 at low pH and high temperature at pH 7.0, are already known to result in different aggregate morphologies [12]. Here we explored whether the two conditions also exposed different aggregation-prone regions (APRs). APRs can be predicted from sequence information, and either assume a fully unfolded protein, or otherwise refine the prediction by factoring solvent exposure of the APR based on structure and dynamics information. The sequence-based predictions are based on either the intrinsic properties of amino acids, or their compatibility with protein structural features in known amyloid fibril structures. Examples of sequence-based predictors include PASTA 2.0 [35], TANGO [36], AGGRESCAN [37], MetAmyl [38], FoldAmyloid [39] and Waltz [40]. As the ability of APRs to trigger aggregation depends upon their solvent accessibility, more recent structure-based predictors consider the three-dimensional structure of the protein and in some cases also their potential modes of partial unfolding. Examples include AGGRESCAN 3D [41], AggScore [42], SAP [43] and Solubis [44]. Here, we want to compare the solvent accessibility of APRs in Fab A33, between our MD simulations at the unfolding conditions and at the reference trajectory. Thus, we used sequence-based APR predictors to determine the APRs in Fab A33, and then determined their solvent accessible surface area (SASA) from the MD simulations, for relative comparisons.

We used four sequence-based predictors to determine the APRs in Fab A33, PASTA 2.0, TANGO, AGGRESCAN and MetAmyl. APRs were selected when three out of the four predictors identified an aggregation-prone sequence (Fig S9A). Seven APRs were found, namely residues 31-36, 47-51, 114-118 and 129-139 in the light chain and residues 261-165, 325-329 and 387-402 in the heavy chain. The presence of these APRs was confirmed using Amylpred2 [45], a consensus tool of eleven existing algorithms (Fig S9A). The location of the seven APRs revealed that they are all located in the interior of Fab A33, and thus protected from the solvent (Fig S9B). Exposure of one of these APRs as a result of a conformational change by an environmental stress, has the potential to trigger aggregation. Thus, the SASA of each APR during the simulations was calculated, as well as the difference in solvent accessibility, ΔSASA, between unfolding conditions and the reference simulation (Table 2).

**Table 2.** SASA of the APRs in Fab A33 during simulations and SASA differences between unfolding simulations and the reference simulation. Solvent accessible surface area of the seven aggregation-prone regions in Fab A33 during all simulations, and relative differences (ΔSASA) between the unfolding trajectories (pH 3.5 and 4.5 at 300 K, for low pH; pH 7.0 at 340 K and 380 K, for high temperature) and the reference trajectory (pH 7.0 and 300 K).

| APR | Fab domain | SASA ($\text{\AA}^2$)<br>pH 7.0<br>300 K | SASA ($\text{\AA}^2$)<br>pH 4.5<br>300 K | ΔSASA ($\text{\AA}^2$)<br>pH(4.5-7.0) | SASA ($\text{\AA}^2$)<br>pH 3.5<br>300 K | ΔSASA ($\text{\AA}^2$)<br>pH(3.5-7.0) | SASA ($\text{\AA}^2$)<br>pH 7.0<br>340 K | ΔSASA ($\text{\AA}^2$)<br>T(340-300 K) | SASA ($\text{\AA}^2$)<br>pH 7.0<br>380 K | ΔSASA ($\text{\AA}^2$)<br>T(380 -<br>300 K) |
| --- | --- | --- | --- | --- | --- | --- | --- | --- | --- | --- |
| 31-36 | $V_L$ | $118 \pm 4$ | $112 \pm 5$ | $-5 \pm 7$ | $115 \pm 16$ | $-3 \pm 17$ | $120 \pm 8$ | $3 \pm 9$ | $106 \pm 4$ | $-12 \pm 6$ |
| 47-51 | $V_L$ | $100 \pm 1$ | $104 \pm 7$ | $4 \pm 13$ | $115 \pm 20$ | $15 \pm 23$ | $101 \pm 6$ | $1 \pm 13$ | $96 \pm 7$ | $-4 \pm 13$ |
| 114-118 | $C_L$ | $125 \pm 4$ | $121 \pm 7$ | $-4 \pm 8$ | $110 \pm 7$ | $-15 \pm 8$ | $112 \pm 2$ | $-13 \pm 5$ | $120 \pm 13$ | $-5 \pm 13$ |
| 129-139 | $C_L$ | $152 \pm 8$ | $147 \pm 11$ | $-5 \pm 13$ | $151 \pm 12$ | $0 \pm 14$ | $165 \pm 10$ | $13 \pm 13$ | $172 \pm 14$ | $20 \pm 16$ |
| 261-265 | V<sub>H</sub> | 14 ± 2 | 13 ± 0 | -1 ± 2 | 13 ± 1 | -1 ± 3 | 19 ± 1 | 5 ± 3 | 29 ± 5 | 15 ± 6 |
| 325-329 | V<sub>H</sub> | 122 ± 16 | 115 ± 14 | -7 ± 21 | 119 ± 7 | -3 ± 17 | 88 ± 10 | -34 ± 19 | 120 ± 22 | -1 ± 27 |
| 387-402 | C<sub>H</sub>1 | 552 ± 21 | 547 ± 25 | -5 ± 33 | 609 ± 12 | 57 ± 25 | 544 ± 7 | -9 ± 22 | 489 ± 48 | -15 ± 21 |

At low pH, only one APR (residues 387-402), was found to increase its solvent accessibility significantly at pH 3.5, with an increase of 57 ± 25 Å^2^ (10 % increase) (Table 2), consistent with previous experimental findings [14]. This APR is located in the C_H_1 domain and its exposure can be explained by the C_L_ domain displacement observed at low pH (Fig 8A). At high temperature, two APRs were found to increase their solvent accessibility, APR 261-265 located in V_H_ and APR 129-139 located in C_L_ (Fig 8B). APR 261-265 increased its SASA 15 ± 6 Å^2^ (107 % increase) and APR 129-139 increased its SASA 20 ± 16 Å^2^ (13 % increase) (Table 2). The location of these APRs agrees with the domains found to unfold previously at high temperature. Notably, the APRs exposed at low pH and high temperature are different, suggesting the potential to follow different aggregation mechanisms depending on the stress applied.

**Fig 8.**
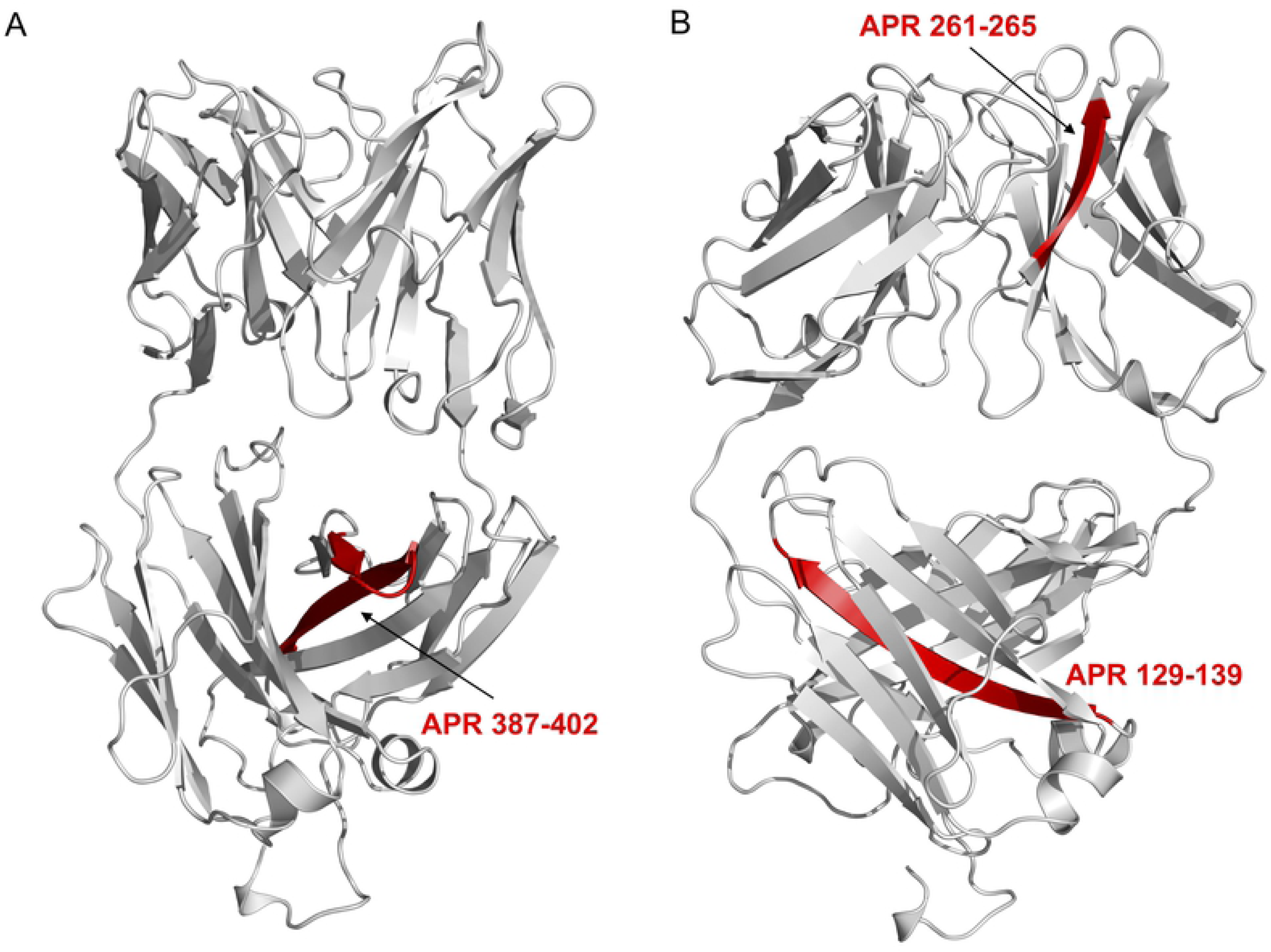
Fab A33 predicted APRs that increase its solvent accessibility at low pH and high temperature. A) APR 387-402 increases its SASA at pH 3.5 and B) APR 261-265 and APR 129-139 increase its SASA at 380 K (Table 1). All mapped in red in Fab A33 homology structure.

## Discussion

Antibody-based products are the main class of approved biopharmaceuticals, due to their high target specificity [1]. However, there are many barriers to their successful development into therapeutics, with protein aggregation being perhaps the most common and challenging to prevent [5]. There is a need to identify potential instabilities of therapeutic proteins during their early development, particularly against stresses that they will encounter during manufacture, storage and delivery. This would allow their early elimination from further development, or otherwise rational mutagenesis into more stable products. In this context, we have elucidated the first unfolding events that take place on a humanized Fab A33 using atomistic MD simulations, and compared these to predictions of potentially stabilising mutations using computational tools.

Our simulations showed that contacts at the interface between domains (V_L_-V_H_ and C_L_-C_H_1) were lost before individual domains unfolded. Interfacial contacts in the constant domain were the least stable, and were lost very quickly during the simulations under both stresses of low pH and high temperature. In line with these results, FoldX and Rosetta agreed that the residues that can be stabilized the most, are located in the constant domain interface. Further validation was provided by packing density calculations, which revealed that the residues identified by the stability predictors, were under packed relative to the other residues located in the interface between domains. Based on these findings, we speculate that improvement of Fab A33 stability should be targeted to the constant domain interface. Only one of the top mutations suggested by FoldX and Rosetta, N137I was found to be present in our analysis of natural variation within existing Fab sequences. However, there was significant scope for improvement through mutating the interfacial residues S176, N137, S397, T180, and S395, to the suggested hydrophobic residues.

The further goal could be to improve the stability of the individual domains. The C_L_ domain was found to unfold at both, low pH and high temperature. Salt bridge analyses identified two key salt bridges at the heart of this domain unfolding at low pH, Glu165-Lys103 and Glu195-Lys149. Glu165-Lys103 bridges the C_L_ domain to the V_L_ domain, and Glu195-Lys149 is located in outer β-strands of the C_L_ domain, bridging β-strands C and F. FoldX and Rosetta also identified stabilizing mutations in the C_L_ domain. To stabilize the interaction between the C_L_ and V_L_ domain, hydrophobic mutants of S12 and K103 were predicted. Interestingly, the mutation S12 to Tyr is found naturally. In the C_L_ domain, S159 was identified, which interacts with an outer β-strand, suggesting this interaction can also be improved. Lastly, the V_H_ domain was also found to unfold at high temperature. The only mutation identified in this domain is S267, to a Pro, which notably is found naturally. Overall, the results found with MD simulations and stabilizing software predictors strongly agree in the domains of Fab A33 that can be stabilized further.

In order to gain insights into the mechanisms by which aggregation might occur, APRs in Fab A33 were identified, and their solvent accessibilities were compared. All APRs in Fab A33 are located in the interior of the protein, however, at low pH and high temperature the SASA of certain APRs increased. Notably, different APRs were exposed under both stresses, suggesting that different aggregation mechanisms occur under each stress. This result stresses the importance of identifying the stability of a protein under the different stresses it might encounter. Taken together, this work provides insights into the stability and robustness of the therapeutically relevant Fab A33, and offers a path to the engineering and design of a more aggregation resistant antibody fragment.

## Materials and Methods

### Fab A33 homology model

The homology model of wild-type Fab A33 was built from the crystal structure of human germline antibody 5-51/O12 (PDB ID: 4KMT) and the amino-acid sequence of Fab A33 (Fig. S1) [46,47]. The C226S heavy-chain variant was used to avoid the formation of linked Fab dimers. We used the Rosetta method “minirosetta”, as detailed in previous works [48].

### Molecular dynamics simulations

Molecular dynamic (MD) simulations on the Fab A33 homology model were conducted in Gromacs v5.0 [49]. MD simulations were carried out at neutral pH and room temperature (pH 7.0 and 300 K) and under two stresses, low pH (pH 3.5 and pH 4.5 at 300 K) and high temperature (pH 7.0 at 340 K and 380 K). Many high temperature simulations are performed at relatively high temperatures (e.g. 500 K), to achieve complete denaturation of the protein, however, in this case, we aim to partially unfold Fab A33 and detect the regions prone to early unfolding. Simulations were carried out using the OPLS-AA/L all-atom force field [50]. The Fab PDB file was first converted to a topology file with its five (four intra-chain and one inter-chain) disulfide bonds retained. The protonation state of each residue was entered manually, and these were determined at each pH using the PDB2PQR server, which performed the pKa calculations by PropKa [51]. This gave the following total charges: +9 (pH 7.0), +18 (pH 4.5) and +35 (pH 3.5). The Fab A33 structure was centered in a cubic box with a layer of water up to at least 10.0 Å from the protein surface. The box was solvated with SPC/E water molecules, Cl-added to neutralize the net charges, and NaCl added to an ionic strength of 50 mM for all simulations. The system was energy minimized using the steepest descent algorithm (2000 steps) followed by the conjugate gradient method (5000 steps). The solvent and ions around the protein were equilibrated in position-restricted simulations for 100 ps under NVT ensemble to stabilize at the specified temperature, and then at 100 ps under NPT ensemble to stabilize at atmospheric pressure. Lastly, MD simulations were carried out for 50 ns in triplicates under the five conditions (pH 7.0 and 300 K; pH 4.5 and 300 K; pH 3.5 and 300 K; pH 7.0 and 340 K; pH 7.0 and 380 K). Jobs were submitted to the UCL Legion High Performance Computing Facility. The time step of the simulations was set to 2 fs and trajectories were saved every 10 ps.

### Analysis of MD trajectories

MD trajectories were saved reduced, every 0.2 ns (total of 250 frames). Interface contacts over simulation time were calculated using the native contacts extension of the visual molecular dynamics (VMD) program [52]. A cutoff distance of 4 Å was used in the calculations. Variable domain contacts (V_L_-V_H_) were calculated between residues 1-108 (V_L_) and 215-334 (V_H_). Constant domain contacts (C_L_-C_H_1) were calculated between residues 109-214 (C_L_) and 335-442 (C_H_1). RMSD of individual domains during the simulations were calculated using the RMSD trajectory tool in VMD. All the structures of the trajectory were first aligned and the RMSD was calculated (no hydrogens included). Domains were V_L_ (1-108), V_H_ (215-334), C_L_ (109 to 214) and C_H_1 (335 to 429). Averages and SEM of three independent repeats are shown. Structural alignments of the last 30 ns of the trajectories were also performed using VMD. Secondary structure (SS) assignments of each residue along the trajectory were done using the DSSP module [53,54]. To analyze the loss in β-strand structure, we monitored the percentage of β-sheet SS per residue. These values were summed for each of the 32 β-strands in Fab A33 and differences were calculated between the unfolding simulations and the reference simulations (pH 7 and 300 K). Lastly, salt bridges were calculated along the trajectories using VMD and a cutoff distance between O and N groups of 3.2 Å. From these, the occurrence (%) of each salt bridge during the simulation was calculated, and averaged for the three independent repeats at each condition.

### Mutational study and **ΔΔ**G calculations by FoldX and Rosetta

The effect of mutations on the stability of Fab A33 was studied using FoldX (foldx.crg.es) [26] and the Rosetta method “ddg_monomer” (www. rosettacommons.org) [27]. Both tools predicted the difference in folding free energy, ΔΔG, between the protein carrying a point mutation and the wildtype. Each of the 442 residues in the Fab A33 were mutated to the other 19 possibilities, totaling 8398 single mutants. FoldX was used as a plugin in the graphical interface YASARA [55]. The “Repair” command was used first to energy minimize the homology model of Fab A33, by rearranging the amino acid side chains. Next, the “BuildModel” command was used to introduce the point mutations, optimize the structure of the new protein variant, and calculate the stability change upon mutation. The Rosetta “ddg_monomer” method was used, where an example of mutation and option files, listing the parameters of the executable, can be seen in previous work [48]. Jobs were submitted to the UCL Legion High-Performance Computing Facility.

### Packing Density

Occluded surface (OS) program was used to calculate the atomic packing of Fab A33 [30,31]. The occluded surface packing (OSP) values are useful for identifying regions of loose packing in a protein. OSP value for each residue are calculated from the collection of extended normals (ray-lengths) that extend outward from the molecular surface until they intersect neighboring van der Waals surface. Analysis of these normals, their respective lengths and the surface area involved in the interaction, defines the packing of each atom in the protein.

### Sequence Entropy of Fab sequences

Fab sequences were retrieved from the Protein Data Bank (PDB) [32], totaling one hundred light chains and one hundred heavy chains. For light chains, kappa (κ) and lambda (λ) chains were included. For κ light chains, λ light chains and heavy chains, sequences from the species human and mouse were used. Sequence alignment and calculation of the sequence entropy for each residue were calculated using Bioedit [33]. The maximum entropy for 21 possible amino acids (including stop codon) is 3.04 and zero represents a fully conserved residue.

### Aggregation-prone regions (APR) predictions

Aggregation prone regions (APR) of Fab A33 were predicted using PASTA 2.0 [35], TANGO [36], AGGRESCAN [37] and MetAmyl [38], using the protein sequence as input. The regions in which three out of the four software identified an APR were selected, resulting in seven APRs. Amylpred2 consensus tool was used to confirm the presence of these APRs [45]. To calculate the solvent accessible surface area (SASA) of each APR during the trajectories, the average area per residue over the trajectory was calculated first, using gromacs analysis tool “sasa”, then summed for each APR.

## ACKNOWLEDGMENTS

We thank the Engineering and Physical Sciences Research Council (EPSRC) Centre for Doctoral Training in Emergent Macromolecular Therapies (EP/L015218/1) (N.C.C.), the EPSRC Future Targeted Healthcare Manufacturing Hub (EP/P006485/1, EP/I033270/1) (N.C.), and EPSRC EP/N025105/1 (C.Z.).

## Supporting information captions

**Fig S1. Fab A33 sequence.** Fab A33 amino acid sequence separated by domains (V_L_, C_L_, V_H_, C_H_1 and hinge region). The six CDRs in the V_L_ and V_H_ domains are highlighted in red.

**Table S1.** Residues located in the interface between light and heavy chains in Fab A33.

**Fig S2. All protein RMSD and radius of gyration (R_g_) with simulation time.** A, B) RMSD of the whole protein with simulation time for different A) pHs and B) temperatures, values as labelled. In all cases, the average of three independent simulations is shown with the SEM as error. C, D) Radius of gyration (R_g_) of Fab A33 with simulation time for different A) pHs and B) temperatures, values as labelled. In all cases, the average of three independent simulations is shown with the SEM as error.

**Fig S3. Secondary structure (SS) of each residue in Fab A33 with simulation time, calculated using DSSP**. Representative SS evolution are shown for A) pH 7.0 and 300K, B) pH 3.5 and 300K and C) pH 7.0 and 380 K, secondary structure type as indicated in the legend.

**Fig S4. Loss of secondary structure for each of the 32 β-strands of Fab A33.** A, B) Strand order shown by lettering (A-G) for variable and constant domains, respectively. C, D) Percentage increase/decrease in β-strand secondary structure for each strand in Fab during the simulations, respect to pH 7.0 and 300 K, for: C) pH 4.5 and 300 K, D) pH 7.0 and 340 K. Error bars are the same and equal for positive and negative values.

**Fig S5. Salt Bridge analysis for simulations at pH 4.5 at 300 K, and pH 7.0 at 340 K.** A, B) Salt bridges formed during the simulation time for representative MD simulations at A) pH 4.5 and 300 K, B) pH 7.0 and 340 K. Presence of a salt bridge is indicated in white and absence in black. C, D) List of salt bridges and its occurrence for simulations at A) pH 4.5 and 300 K, B) pH 7.0 and 340 K. Values shown are the average of three independent simulations with their SEM as error. The more persistent salt bridges are highlighted for pH 4.5 (red) and for high temperature 340 K (blue). E, F) The more persistent salt bridges are mapped into the Fab A33 structure.

**Fig S6. Stabilizing mutations predicted by FoldX and Rosetta. Correlation between FoldX and Rosetta predictions.** Mutations predicted by both software to be most stabilizing are shown in magenta and highlighted in a gray square on the bottom left. Mutations predicted only by FoldX to be stabilizing are shown in green and mutations predicted only by Rosetta in yellow.

**Table S2.** List of the most stabilizing mutations identified by FoldX and Rosetta-ddG. **Mutation and ΔΔG of the 25 most stabilizing mutations predicted by FoldX and Rosetta.**

**Fig S7. Normals used to calculate the packing of each atom in Fab A33 using occluded surface software.** To calculate the occluded surface packing (OSP) value for each residue, normals that extend from the surface outward until they intersect a neighboring van der Waals surface are used. The normals used to calculate the OSP value of the inter-domain residues identified by FoldX and Rosetta (S176, N137, S397, T180, and S395), are shown.

**Table S3. Packing indicated by the occluded surface packing (OSP) value of the residues located in β-strands within domain interfaces (V_L_-V_H_ and C_L_-C_H_1) of Fab A33 homology model**. OSP values were calculated using the occluded surface software. *, residues identified by FoldX and Rosetta in the constant domain interface that can be stabilized further.

**Fig S8. Sequence entropy of Fab sequences.** (A) Entropy (Hx) of one hundred Fab light chains, including κ and λ light chains. (B) Entropy (Hx) of one hundred Fab heavy chains. Light and heavy chains are from human and mouse species. Alignment and entropy calculation were done with Bioedit. Variable domains are indicated with a (V) and constant domains with a (C), separated by a vertical line. CDRs are indicated in red.

**Fig S9. Aggregation prone regions (APR) in Fab A33.** A) The normalised aggregation propensity for each residue in Fab A33 was predicted using PASTA 2.0, TANGO, AGGRESCAN and MetAmyl software, each being colour-coded as shown in the legend. Aggregation-prone sequences where three of the four software agreed were selected, and highlighted with an asterisk. Amylpred2 consensus tool was also used to confirm the identification of those APRs, indicated in red on the top. B) Using the native Fab A33 homology model, regions with greater aggregation propensities are shown in red and reduced propensities in blue.

